# Leaky severe combined immunodeficiency in mice lacking non-homologous end joining factors XLF and MRI

**DOI:** 10.1101/2020.03.04.976829

**Authors:** Sergio Castañeda-Zegarra, Qindong Zhang, Amin Alirezaylavasani, Marion Fernandez-Berrocal, Rouan Yao, Valentyn Oksenych

**Author notes:** These authors contributed equally (Sergio Castañeda-Zegarra, Qindong Zhang).

## Abstract

Non-homologous end-joining (NHEJ) is a DNA repair pathway required to detect, process, and ligate DNA double-stranded breaks (DSBs) throughout the cell cycle. The NHEJ pathway is necessary for V(D)J recombination in developing B and T lymphocytes. During NHEJ, Ku70 and Ku80 form a heterodimer that recognizes DSBs and promotes recruitment and function of downstream factors PAXX, MRI, DNA-PKcs, Artemis, XLF, XRCC4, and LIG4. Mutations in several known NHEJ genes result in severe combined immunodeficiency (SCID). Inactivation of *Mri, Paxx* or *Xlf* in mice results in normal or mild phenotype, while combined inactivation of *Xlf/Mri, Xlf/Paxx*, or *Xlf*/*Dna-pkcs* leads to late embryonic lethality. Here, we describe three new mouse models. We demonstrate that deletion of *Trp53* rescues embryonic lethality in mice with combined deficiencies of *Xlf* and *Mri*. Furthermore, *Xlf*^*-/-*^*Mri*^*-/-*^*Trp53*^*+/-*^ and *Xlf*^*-/-*^*Paxx*^*-/-*^*Trp53*^*+/-*^ mice possess reduced body weight, severely reduced mature lymphocyte counts, and accumulation of progenitor B cells. We also report that combined inactivation of *Mri/Paxx* results in live-born mice with modest phenotype, and combined inactivation of *Mri/Dna-pkcs* results in embryonic lethality. Therefore, we conclude that XLF is functionally redundant with MRI and PAXX during lymphocyte development *in vivo*. Moreover, *Mri* genetically interacts with *Dna-pkcs* and *Paxx*.

## 1. Introduction

Non-homologous end-joining (NHEJ) is a DNA repair pathway that recognizes, processes and ligates DNA double-stranded breaks (DSB) throughout the cell cycle. NHEJ is required for lymphocyte development; in particular, to repair DSBs induced by the recombination activating genes (RAG) 1 and 2 in developing B and T lymphocytes, and by activation-induced cytidine deaminase (AID) in mature B cells [1]. NHEJ is initiated when Ku70 and Ku80 (Ku) are recruited to the DSB sites. Ku, together with DNA-dependent protein kinase, catalytic subunit (DNA-PKcs), forms the DNA-PK holoenzyme [2]. Subsequently, the nuclease Artemis is recruited to the DSB sites to process DNA hairpins and overhangs [3]. Finally, DNA ligase IV (LIG4), X-ray repair cross-complementing protein 4 (XRCC4) and XRCC4-like factor (XLF) mediate DNA end ligation. The NHEJ complex is stabilized by a paralogue of XRCC4 and XLF (PAXX) and a modulator of retroviral infection (MRI/CYREN) [4, 5].

Inactivation of *Ku70, Ku80, Dna-pkcs* or *Artemis* results in severe combined immunodeficiency (SCID) characterized by lack of mature B and T lymphocytes [2, 3, 6-8]. Deletion of both alleles of *Xrcc4* [9] or *Lig4* [10] results in late embryonic lethality in mice, which correlates with increased apoptosis in the central nervous system (CNS). Inactivation of *Xlf* (*Cernunnos*) only results in modest immunodeficiency in mice [11-13], while mice lacking *Paxx* [14-17] or *Mri* [5, 18] display no overt phenotype.

The mild phenotype observed in mice lacking XLF could be explained by functional redundancy between XLF and multiple DNA repair factors, including *Ataxia telangiectasia* mutated (ATM), histone H2AX [19], Mediator of DNA Damage Checkpoint 1 (MDC1) [20, 21], p53-binding protein 1 (53BP1) [17, 22], RAG2 [23], DNA-PKcs [20, 24, 25], PAXX [4, 14, 15, 20, 26-28] and MRI [5]. However, combined inactivation of *Xlf* and *Paxx* [4, 14, 15, 20], as well as *Xlf* and *Mri* [5], results in late embryonic lethality in mice, presenting a challenge to the study of B and T lymphocyte development *in vivo*. It has also been shown that both embryonic lethality and increased levels of CNS neuronal apoptosis in mice with deficiency in *Lig4* [9, 10, 29, 30], *Xrcc4* [9, 31], *Xlf* and *Paxx* [20], or *Xlf* and *Dna-pkcs* [24, 25] is p53-dependent. In this study, we rescue synthetic lethality from *Xlf* and *Mri* by inactivating one or two alleles of *Trp53*. We also show that both *Xlf*^*-/-*^*Mri*^*-/-*^*Trp53*^*+/-*^ and *Xlf*^*-/-*^*Paxx*^*-/-*^*Trp53*^*+/-*^ mice possess a leaky SCID phenotype with severely reduced mature B and T lymphocyte counts in the spleen, low mature T cell counts in the thymus, and accumulated progenitor B cells in the bone marrow. Finally, we demonstrate that MRI is functionally redundant with DNA-PKcs and PAXX.

## 2. Materials and Methods

### 2.1. Mice

All experiments involving mice were performed according to the protocols approved by the Comparative Medicine Core Facility (CoMed) at the Norwegian University of Science and Technology (NTNU, Trondheim, Norway). *Xlf*^*+/-*^ [11] and *Dna-pkcs*^*+/-*^ [2] mice were imported from the laboratory of Professor Frederick W. Alt at Harvard Medical School. *Trp53*^*+/-*^ mice [32] were imported from Jackson Laboratories. *Paxx*^*+/-*^ [16] and *Mri*^*+/-*^ [18] mice were generated by the Oksenych group and described previously.

### 2.2. Lymphocyte development

Lymphocyte populations were analyzed by flow cytometry [16, 18, 19, 22]. In summary, cells were isolated from the spleen, thymus, and femur of 5-7-week-old mice and treated with red blood cell lysis buffer Hybri-Max^™^ (Sigma Aldrich, St. Louis, MO, USA; #R7757). The cells were resuspended in PBS (Thermo Scientific, Basingstoke, UK; #BR0014G) containing 5% Fetal bovine serum, FCS (Sigma Life Science, St. Louis, Missouri, United States; #F7524), and counted using a Countess™ II Automated Cell Counter (Invitrogen, Carlsbad, CA, United States; #A27977). Then, the cell suspension was diluted with PBS to get a final cell concentration of 2.5 × 10^7^ cells/mL. Finally, surface markers were labeled with fluorochrome-conjugated antibodies and the cell populations were analyzed using flow cytometry.

### 2.3. Class switch recombination (CSR)

Spleens were isolated from 5-7-week-old mice and stored in cold PBS. Splenocytes were obtained by mincing the spleens, and naïve B cells were negatively selected using an EasySep Isolation kit (Stemcell™, Cambridge, UK; #19854). Lipopolysaccharide (LPS; 40 μg/mL; Sigma Aldrich, St. Louis, MO, USA; #437627-5MG) and interleukin 4 (IL-4; 20 ng/mL; PeproTech, Stockholm, Sweden; #214-14) were used to induce CSR to IgG1. Expression of IgG1 was analyzed by flow cytometry.

### 2.4. Antibodies

The following antibodies were used for flow cytometric analysis: rat anti-CD4-PE-Cy7 (BD Pharmingen^™^, Allschwil, Switzerland, #552775; 1:100); rat anti-CD8-PE-Cy5 (BD Pharmingen^™^, Allschwil, Switzerland, #553034; 1:100); anti-CD19-PE-Cy7 (Biolegend, San Diego, CA, USA, #115520; 1:100); hamster anti-mouse anti-CD3-FITC (BD Pharmingen^™^, Allschwil, Switzerland, #561827; 1:100); rat anti-mouse anti-CD43-FITC (BD Pharmingen^™^, Allschwil, Switzerland, #561856; 1:100); rat anti-mouse anti-CD45R/B220-APC (BD Pharmingen^™^, Allschwil, Switzerland; #553092; 1:100); rat anti-mouse anti-IgM-PE-Cy7 (BD Pharmingen^™^, Allschwil, Switzerland, #552867; 1:100); rat anti-mouse IgG1-APC (BD Pharmingen^™^, Allschwil, Switzerland; #550874; 1:100). A LIVE/DEAD™ fixable violet dead cell stain kit (ThermoFisher Scientific, Waltham, MA, USA; #L34955; 1:1000) was used to identify dead cells.

### 2.5. Statistics

Statistical analyses were performed using one-way ANOVA, GraphPad Prism 8.0.1.244 (San Diego, CA, USA). In all statistical tests, *p*<0.05 were taken to be significant (\**p*<0.05; \*\**p*<0.01; \*\*\**p*<0.001; \*\*\*\**p*<0.0001).

## 3. Results

### 3.1. Inactivation of Trp53 gene rescued embryonic lethality in mice lacking XLF and MRI

Combined inactivation of *Xlf* and *Mri* has previously been shown to result in synthetic lethality in mice [5]. To generate XLF/MRI deficient mice with altered expression of *Trp53*, we intercrossed an *Mri*^*-/-*^ strain [18] with an *Xlf*^*-/-*^*Trp53*^*+/-*^ [20] strain. Next, we selected and intercrossed triple heterozygous (*Xlf*^*+/-*^*Mri*^*+/-*^*Trp53*^*+/-*^), and later, *Xlf*^*-/-*^*Mri*^*+/-*^*Trp53*^*+/-*^ mice. With PCR screening, we identified *Xlf*^*-/-*^*Mri*^*-/-*^*Trp53*^*+/-*^ (n=11), *Xlf*^*-/-*^*Mri*^*-/-*^*Trp53*^*-/-*^ (n=2), and *Xlf*^*-/-*^*Mri*^*-/-*^*Trp53*^*+/+*^ (n=1) (Figure 1A) among the resulting offspring. Mice lacking both XLF and MRI possessed reduced weight (12 g on average, *p*<0.0001) when compared with gender- and age-matched WT (19 g), *Xlf*^*-/-*^ (19 g) and *Mri*^*-/-*^ (20 g) controls (Figure 1B and 1C). In addition, *Xlf*^*-/-*^*Mri*^*-/-*^*Trp53*^*+/-*^ and *Xlf*^*-/-*^*Mri*^*-/-*^*Trp53*^*-/-*^ mice were viable up to 63 days and died for unknown reasons. We used *Xlf*^*-/-*^*Mri*^*-/-*^*Trp53*^*+/-*^ mice to further characterize the development of B and T lymphocytes *in vivo*.

**Figure 1.**
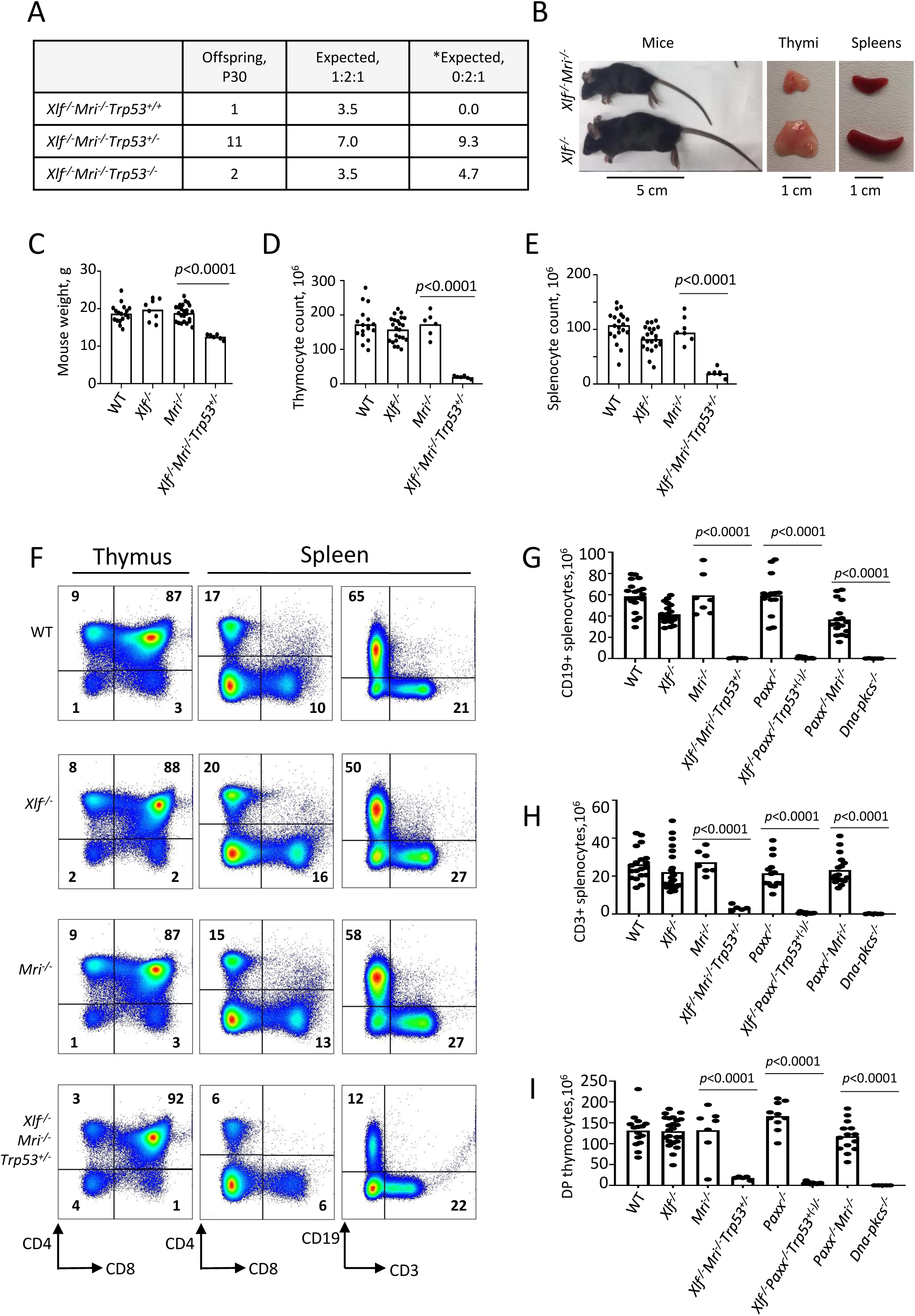
Development of B and T lymphocytes in *Xlf*^*-/-*^*Mri*^*-/-*^*Trp53*^*+/-*^ mice. (A) Number of thirty-day-old mice (P30) of indicated genotypes. *Expected distribution assuming lethality. (B) Comparison of body size, thymi and spleens of XLF/MRI-deficient and XLF-deficient mice of the same age. (C) Weights of WT, *Xlf*^*-/-*^, *Mri*^*-/-*^, *Xlf*^*-/-*^*Mri*^*-/-*^*Trp53*^*+/-*^ mice. (D, E) Number (×10^6^) of thymocytes (D) and splenocytes (E) in WT, *Xlf*^*-/-*^, *Mri*^*-/-*^, *Xlf*^*-/-*^*Mri*^*-/-*^*Trp53*^*+/-*^ mice. (F) Flow cytometric analysis of thymic and splenic T cell subsets and splenic B cells. (G,H,I) Number (×10^6^) of splenic CD19+ B cells (G), splenic CD3+ T cells (H) and thymic CD4+CD8+ double positive (DP) T cells (I) in WT, *Xlf*^*-/-*^, *Mri*^*-/-*^, *Xlf*^*-/-*^*Mri*^*-/-*^*Trp53*^*+/-*^, *Paxx*^*-/-*^, *Xlf*^*-/-*^ *Paxx*^*-/-*^*Trp53*^*+(-)/-*^ and *Paxx*^*-/-*^*Mri*^*-/-*^ mice. *Dna-pkcs*^*-/-*^ mice were used as an immunodeficient control. Comparisons between every two groups were made using one-way ANOVA, GraphPad Prism 8.0.1. *Xlf*^*-/-*^*Paxx*^*-/-*^*Trp53*^*+(-)/-*^ is a combination of *Xlf*^*-/-*^*Paxx*^*-/-*^*Trp53*^*+/-*^ and *Xlf*^*-/-*^*Paxx*^*-/-*^*Trp53*^*-/-*^. Not shown in the graph for (G): WT vs *Paxx*^*-/-*^*Mri*^*-/-*^, *p*<0.0001 (****), *Paxx*^*-/-*^ vs *Paxx*^*-/-*^*Mri*^*-/-*^, *p*<0.0001 (****), *Mri*^*-/-*^ vs *Paxx*^*-/-*^*Mri*^*-/-*^, *p*<0.0025 (**), *Xlf*^*-/-*^ vs *Paxx*^*-/-*^*Mri*^*-/-*^, *p*=0.9270 (n.s), *Xlf*^*-/-*^*Mri*^*-/-*^*Trp53*^*+/-*^ vs *Paxx*^*-/-*^*Mri*^*-/-*^, *p*<0.0001 (****), and *Xlf*^*-/-*^*Paxx*^*-/-*^*Trp53*^*+(-)/-*^ vs *Paxx*^*-/-*^*Mri*^*-/-*^, *p*<0.0001 (****).

### 3.2. Leaky SCID in Xlf^-/-^Mri^-/-^Trp53^+/-^ mice

To determine the roles of XLF and MRI in lymphocyte development *in vivo*, we isolated the thymus, spleen, and femur from *Xlf*^*-/-*^*Mri*^*-/-*^*Trp53*^*+/-*^ mice, as well as from *Xlf*^*-/-*^, *Mri*^*-/-*^, *Trp53*^*+/-*^ and WT controls. Combined deficiency for XLF and MRI resulted in a 3-fold reduction in thymus size (32 mg on average, *p*<0.0001) and a 9-fold reduction in thymocyte count (1.9×10^7^, *p*<0.0001) when compared to single deficient or WT controls (Figure 1D). Similarly, both average spleen weight (22 mg, *p*<0.0001) and splenocyte count (2.0×10^7^, *p*<0.0001) in *Xlf*^*-/-*^*Mri*^*-/-*^*Trp53*^*+/-*^ mice decreased approximately 4-5 fold when compared with WT and single deficient controls (Figure 1E). The reduced number of splenocytes in XLF/MRI double-deficient mice could be explained by decreased populations of B and T lymphocytes observed in the *Xlf*^*-/-*^*Mri*^*-/-*^*Trp53*^*+/-*^ mice (Figure 1F-H). Specifically, CD3+ T cells were reduced 6-fold (*p*<0.0001), while CD19+ B cells were reduced 50-fold (*p*<0.0001) when compared with single deficient and WT controls (Figure 1F-H). Likewise, counts of CD4+ and CD8+ T cells in the spleen, were all dramatically reduced when compared with single deficient and WT controls (about 4-fold, *p*<0.0001; Figure 1F, 1H) as well as counts of CD4+, CD8+ and CD4+CD8+ T cells in the thymus (Figure 1F,I). From these observations, we conclude that XLF and MRI are functionally redundant during B and T lymphocytes development in mice.

### 3.3. Leaky SCID in mice lacking XLF and PAXX

Combined inactivation of XLF and PAXX has been shown to result in embryonic lethality in mice [4, 14, 15, 20]. To determine the impact of XLF and PAXX on B and T cell development *in vivo*, we rescued the synthetic lethality by inactivating one allele of *Trp53*, as described previously [20]. We did not detect any direct influence of altered *Trp53* genotype on lymphocyte development. The resulting *Xlf*^*-/-*^*Paxx*^*-/-*^*Trp53*^*+/-*^ and *Xlf*^*-/-*^*Paxx*^*-/-*^*Trp53*^*-/-*^ mice possess 30-to 40-fold reduced thymocyte count (4.0×10^6^, *p*<0.0001) when compared to WT (1.3×10^8^), *Xlf*^*-/-*^ (1.4×10^8^) and *Paxx*^*-/-*^ (1.7×10^8^) mice. This is reflected in decreased levels of double-positive CD4+CD8+ cells, as well as decreased levels of single-positive CD4+ and CD8+ T cells (Figure 1, Supplementary Figure 1). Spleen development was dramatically affected in mice lacking XLF and PAXX compared to WT and single-deficient controls, due to the lack of B cells and decreased T cell count (Figure 1, Supplementary Figure 1). When compared with the WT and single knockout controls, *Xlf*^*-/-*^*Paxx*^*-/-*^*Trp53*^*+/-*^ and *Xlf*^*-/-*^*Paxx*^*-/-*^*Trp53*^*-/-*^ mice had a 100- to 600-fold reduction in CD19+ B splenocyte count (0.7×10^6^, *p*<0.0001) and a 50- to 90-fold reduction in CD3+ splenocyte count (to 0.5×10^6^) (Figure 1F-H and Supplementary Figure 1). From these results, we concluded that XLF and PAXX are functionally redundant during the B and T lymphocyte development *in vivo*.

### 3.4. Early B cell development is abrogated in mice lacking XLF and MRI, or XLF and PAXX

Reduced counts and proportions of mature B lymphocytes in *Xlf*^*-/-*^*Mri*^*-/-*^*Trp53*^*+/-*^ mice suggest a blockage in B cell development in the bone marrow. To investigate this further, we isolated the bone marrow cells from femora of mice lacking XLF, MRI or both XLF/MRI, and analyzed the proportions of B220+CD43+IgM-progenitor B cells and B220+CD43-IgM+ immature and mature B cells. We detected only background levels of B220+CD43-IgM+ B cells in bone marrows isolated from *Xlf*^*-/-*^*Mri*^*-/-*^*Trp53*^*+/-*^ mice (Figure 2A, 2B). However, these mice exhibited a 2- to 3-fold higher proportion of pro-B cells when compared with WT, *Xlf*^*-/-*^ and *Mri*^*-/-*^ controls (Figure 2A, 2C). Similarly, *Xlf*^*-/-*^*Paxx*^*-/-*^*Trp53*^*+/-*^ and *Xlf*^*- /-*^*Paxx*^*-/-*^*Trp53*^*-/-*^ mice also possess background levels of IgM+ B cells (*p*<0.0001; Figure 2A,B) while having 3- to 4-fold higher proportion of pro-B cells when compared with WT, *Xlf*^*-/-*^ and *Paxx*^*-/-*^ controls (*p*<0.0001; Figure 2A,C). Therefore, we conclude that B cell development is blocked at the pro-B cell stage of *Xlf*^*-/-*^*Mri*^*-/-*^*Trp53*^*+/-*^ and *Xlf*^*-/-*^*Paxx*^*-/-*^*Trp53*^*+/-*^ mice.

**Figure 2.**
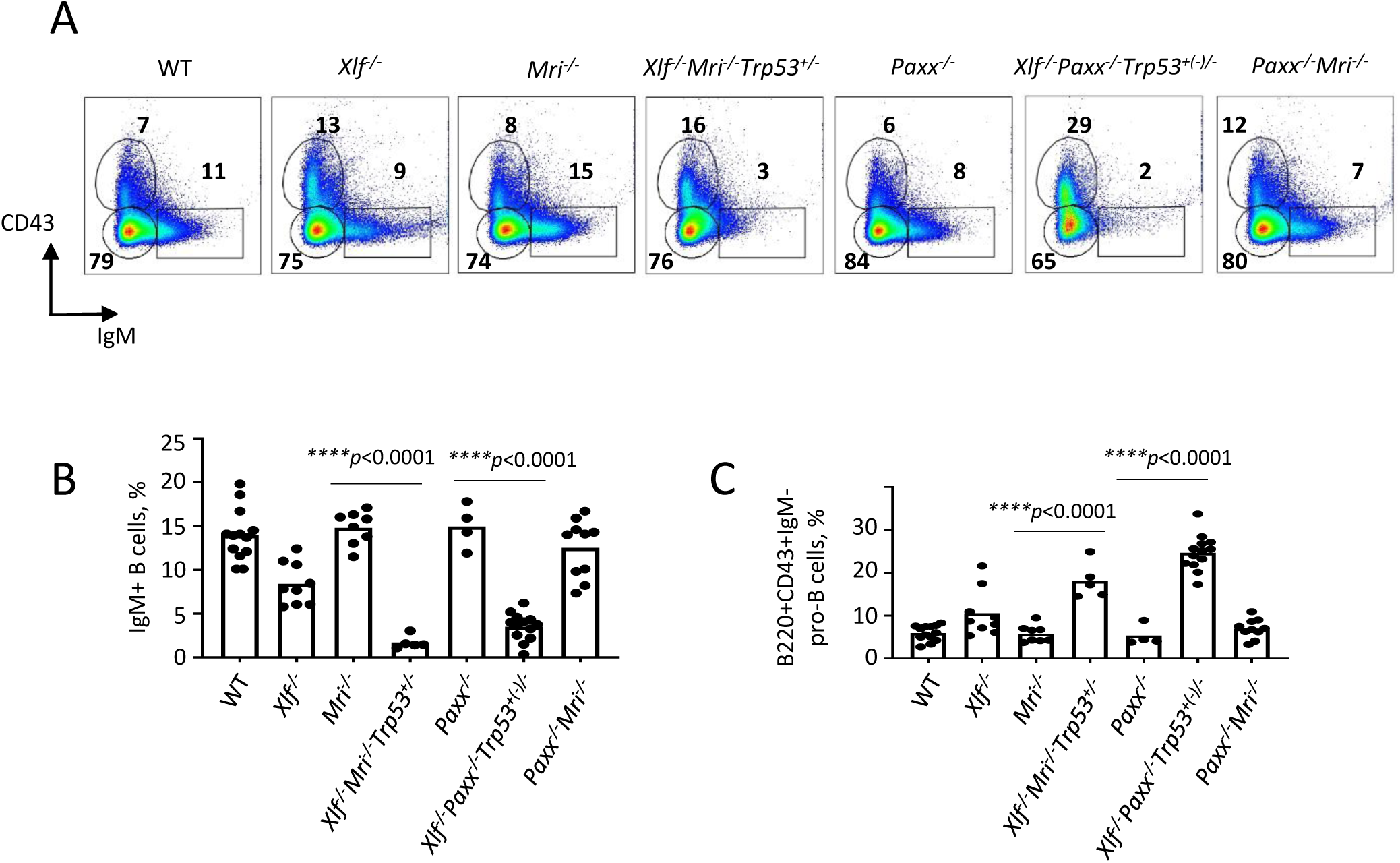
Development of B cells is abrogated in bone marrow of *Xlf*^*-/-*^*Mri*^*-/-*^*Trp53*^*+/-*^ and *Xlf*^*-/-*^*Paxx*^*-/-*^ *Trp53*^*+/-*^ mice. (A) Flow cytometric analysis of developing B cells. Upper left boxes mark B220+CD43+IgM-progenitor B cell populations, and lower right boxes mark the B220+CD43-IgM+ B cells. (B, C) Frequencies (%) of B220+CD43-IgM+ B cells (B) and B220+CD43+IgM-progenitor B cells (C) in WT, *Xlf*^*-/-*^, *Mri*^*-/-*^, *Xlf*^*-/-*^*Mri*^*-/-*^*Trp53*^*+/-*^, *Paxx*^*-/-*^, *Xlf*^*-/-*^*Paxx*^*-/-*^*Trp53*^*+(-)/-*^ and *Paxx*^*-/-*^*Mri*^*-/-*^ mice. Comparisons between groups were made using one-way ANOVA, GraphPad Prism 8.0.1. *Xlf*^*-/-*^*Paxx*^*-/-*^ *Trp53*^*+(-)/-*^ is a combination of *Xlf*^*-/-*^*Paxx*^*-/-*^*Trp53*^*+/-*^ and *Xlf*^*-/-*^*Paxx*^*-/-*^*Trp53*^*-/-*^.

### 3.5. *Paxx*^*-/-*^*Mri*^*-/-*^ *mice* possess a modest phenotype

Both PAXX and MRI are NHEJ factors that are functionally redundant with XLF in mice. Combined inactivation of *Paxx* and *Xlf* [4, 14, 15, 20], or *Mri* and *Xlf* ([5]; this study) results in synthetic lethality in mice, as well as in abrogated V(D)J recombination in vAbl pre-B cells [4, 5, 14, 15, 27]. To determine if *Paxx* genetically interacts with *Mri*, we intercrossed mice that are heterozygous or null for both genes (such as *Paxx*^*-/-*^*Mri*^*+/-*^ and *Paxx*^*+/-*^*Mri*^*+/-*^). We found that resulting *Paxx*^*-/-*^*Mri*^*-/-*^ mice are live-born, fertile, and are similar in size to WT littermates (17 g, *p*>0.9999) (Figure 3A and 3B). Specifically, we observe that *Paxx*^*-/-*^*Mri*^*-/-*^ mice have normal thymocyte and splenocyte counts. Furthermore, *Paxx*^*-/-*^*Mri*^*-/-*^ mice underwent normal T cell development that was indistinguishable from the WT, *Paxx*^*-/-*^, and *Mri*^*-/-*^ controls (Figure 1H, 1I and 3C). However, *Paxx*^*-/-*^*Mri*^*-/-*^ mice had reduced CD19+ B cell counts (Figure 1G) when were compared to WT, *Paxx*^*-/-*^ and *Mri*^*-/-*^ controls (*p*<0.0025). Moreover, CD19+ B cell counts were similar in *Paxx*^*-/-*^*Mri*^*-/-*^ and *Xlf*^*-/-*^ mice (p>0.9270), suggesting that combined depletion of PAXX and MRI has modest phenotype similar to the one in XLF-deficient mice. CSR to IgG1 was performed in order to determine if DNA repair-dependent immunoglobulin production is affected in mature B cells lacking PAXX and MRI [16, 18]. *Paxx* inactivation did not affect Ig switch to IgG1 in MRI-deficient B cells (Figure 3D and 3E). The quantity of IgG1+ cells after CSR stimulation was similar between *Paxx*^*-/-*^*Mri*^*-/-*^ and *Mri*^*-/-*^ naïve B cells (p>0.73). From this, we can conclude that there is a genetic interaction between *Paxx* and *Mri in vivo*, and it is only detected in B cells.

**Figure 3.**
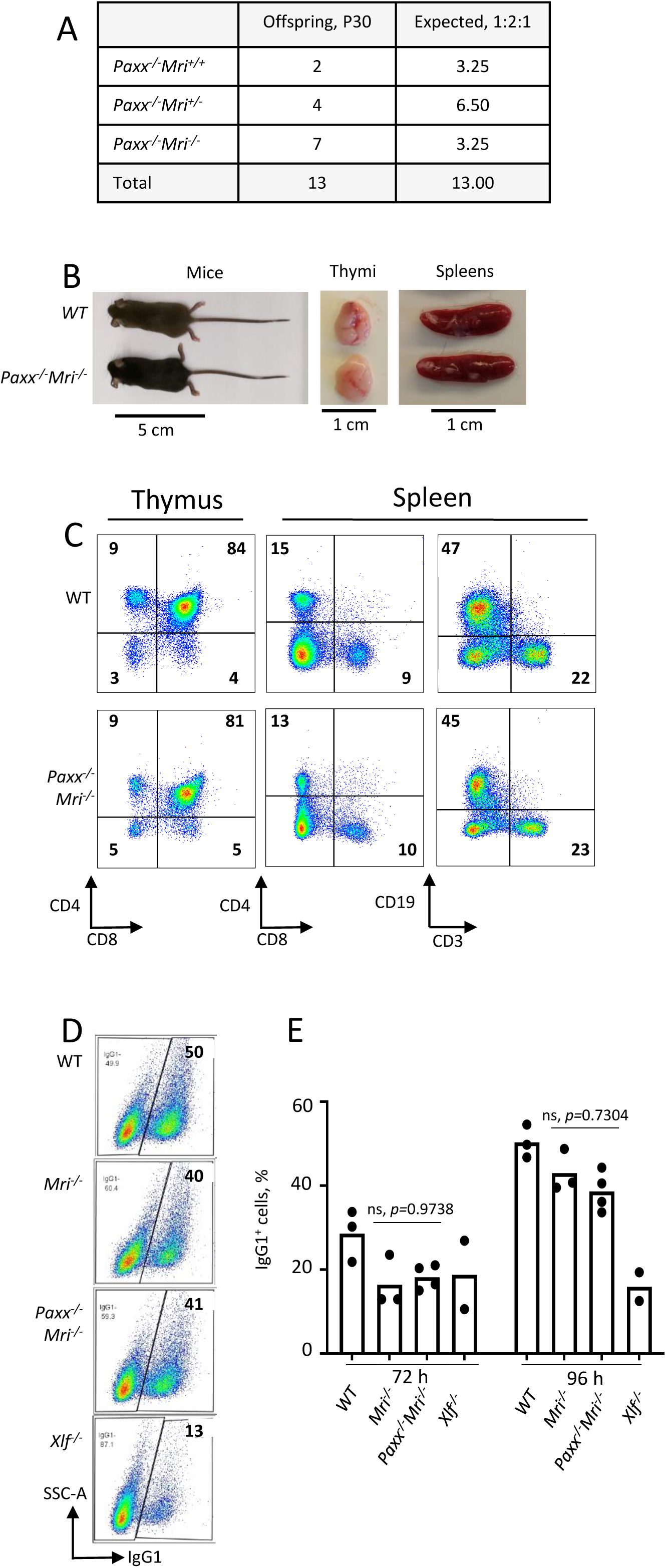
Development of B and T cells in *Paxx*^*-/-*^*Mri*^*-/-*^ mice. (A) Number of thirty-day-old mice (P30) of indicated genotypes. Parents were *Paxx*^*+/-*^*Mri*^*+/-*^ and *Paxx*^*-/-*^*Mri*^*+/-*^. (B) Example of thirty-day-old *Paxx*^*- /-*^*Mri*^*-/-*^ and WT male littermates with their respective thymi and spleens. (C) Example of flow cytometry analyzes of B and T cells in *Paxx*^*-/-*^*Mri*^*-/-*^ and WT mice. (D,E) Class switching analyzes of *in vitro* activated naïve B cells of indicated genotypes.

### 3.6. Synthetic lethality between Mri and Dna-pkcs in mice

Both MRI and DNA-PKcs are functionally redundant with XLF in mouse development [5, 24]. Combined inactivation of *Paxx* and *Mri* (this study), or *Paxx* and *Dna-pkcs* [20] genes results in live-born mice that are indistinguishable from single deficient controls. To determine if *Mri* genetically interacts with *Dna-pkcs*, we crossed *Mri*^*+/-*^ and *Dna-pkcs*^*+/-*^ mouse strains, then intercrossed the double-heterozygous *Mri*^*+/-*^*Dna-pkcs*^*+/-*^, and then *Mri*^*-/-*^*Dna-pkcs*^*+/-*^ mice (Figure 4A). We identified 12 *Mri*^*-/-*^*Dna-pkcs*^*+/+*^ and 12 *Mri*^*-/-*^*Dna-pkcs*^*+/-*^, but no *Mri*^*-/-*^*Dna-pkcs*^*-/-*^ mice (out of 6 expected). To determine if double-deficient *Mri*^*-/-*^*Dna-pkcs*^*-/-*^ embryos are present at day E14.5, we intercrossed *Mri*^*- /-*^*Dna-pkcs*^*+/-*^ mice, extracted and genotyped the embryos (Figure 4B). We identified two *Mri*^*-/-*^*Dna-pkcs*^*-/-*^ mice at E14.5 (63mg), which were about 40% lighter than *Mri*^*-/-*^ littermates (108mg) (Figure 4C and 4D). A Chi-Square test (χ^2^) was performed to determine if the embryonic distribution data fits the mendelian ratio of 1:2:1 that is expected from *Mri*^*-/-*^*Dna-pkcs*^*+/-*^ parents. With DF=2 and χ^2^=1.8, the corresponding p-value lies within the range 0.25<p<0.5. This affirms that our data fit the expected 1:2:1 distribution and suggests that *Mri*^*-/-*^*Dna-pkcs*^*-/-*^ is synthetic lethal. Therefore, we can conclude that there is genetic interaction between *Mri* and *Dna-pkcs in vivo*.

**Figure 4.**
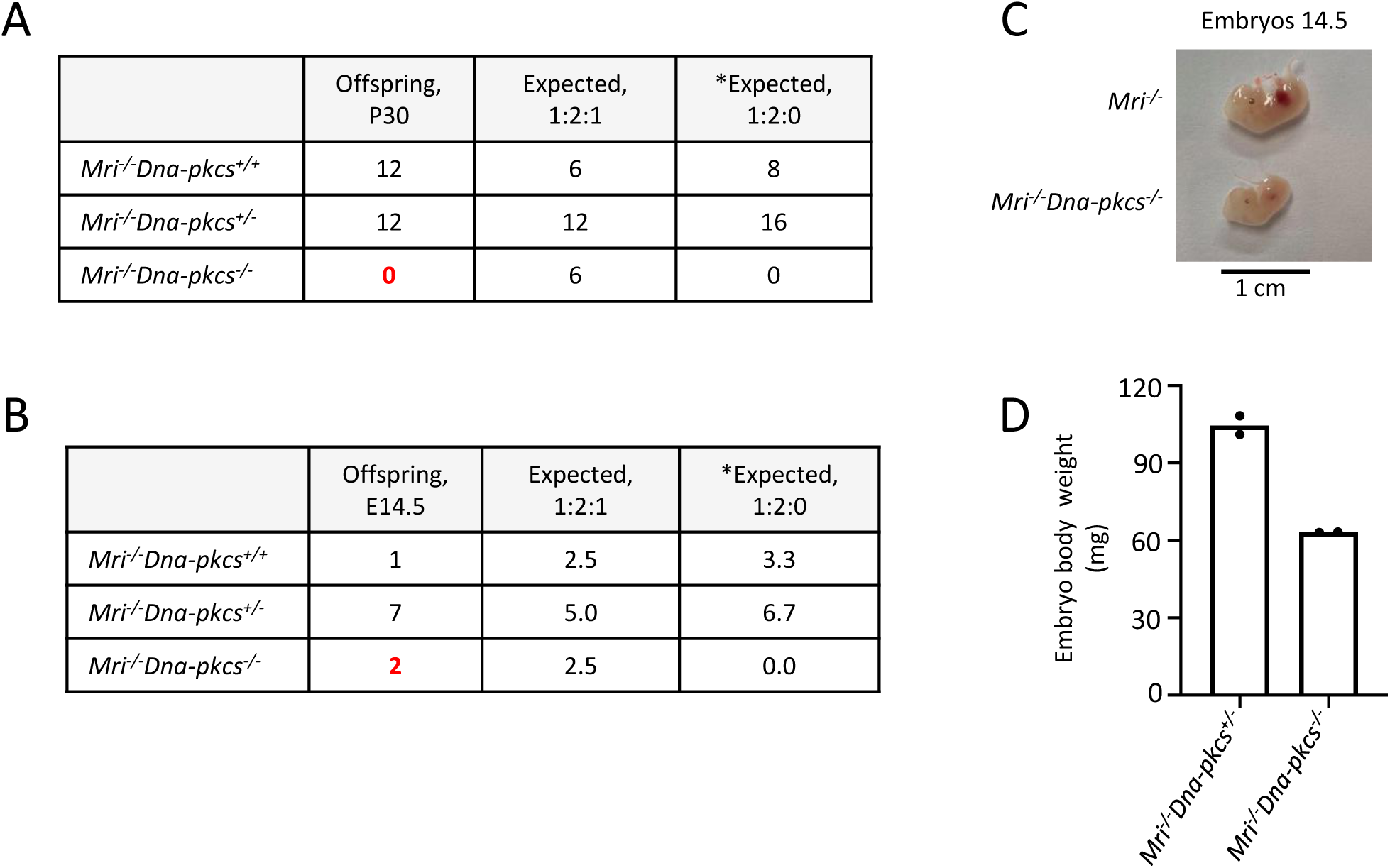
Genetic interaction between *Mri* and *Dna-pkcs in vivo*. (A) No live-born *Mri*^*-/-*^*Dna-pkcs*^*-/-*^ mice were detected. (B,C) *Mri*^*-/-*^*Dna-pkcs*^*-/-*^ embryos were detected at day E14.5. (D) Body weight in milligrams (mg) from two E14.5 *Mri*^*-/-*^*Dna-pkcs*^*-/-*^ and *Mri*^*-/-*^*Dna-pkcs*^*+/-*^ embryos from the same litter. The mendelian ratio 1:2:1 in embryos was verified by the Chi-Square test (χ^2^). The χ^2^ was 1.8 and its corresponding probability was between 25 and 50%. *Expected distribution assuming lethality.

## 4. Discussion

Recent findings by our and other research groups suggest that MRI forms heterogeneous complexes involving PAXX or XLF, which function during DNA DSB repair by NHEJ [5]. Furthermore, genetic inactivation of *Xlf* [11], *Paxx* [4, 14-16], or *Mri* [5, 18] in mice leads to development of modest or no detectable phenotype. However, combined inactivation of *Xlf* and *Mri* [5] or *Xlf* and *Paxx* [4, 14, 15] results in embryonic lethality, which correlates with increased levels of neuronal apoptosis in the CNS (Figure 5). Here, we show that synthetic lethality produced by combined inactivation of *Xlf* and *Mri* can be rescued by altered *Trp53* expression, similar to our previous *Xlf*^*-/-*^*Paxx*^*-/-*^*Trp53*^*+(-)/-*^ [20] mouse model. Furthermore, we have developed and presented here *Paxx*^*-/-*^*Mri*^*-/-*^ and *Mri*^*-/-*^*Dna-pkcs*^*-/-*^ double deficient models.

**Figure 5.**
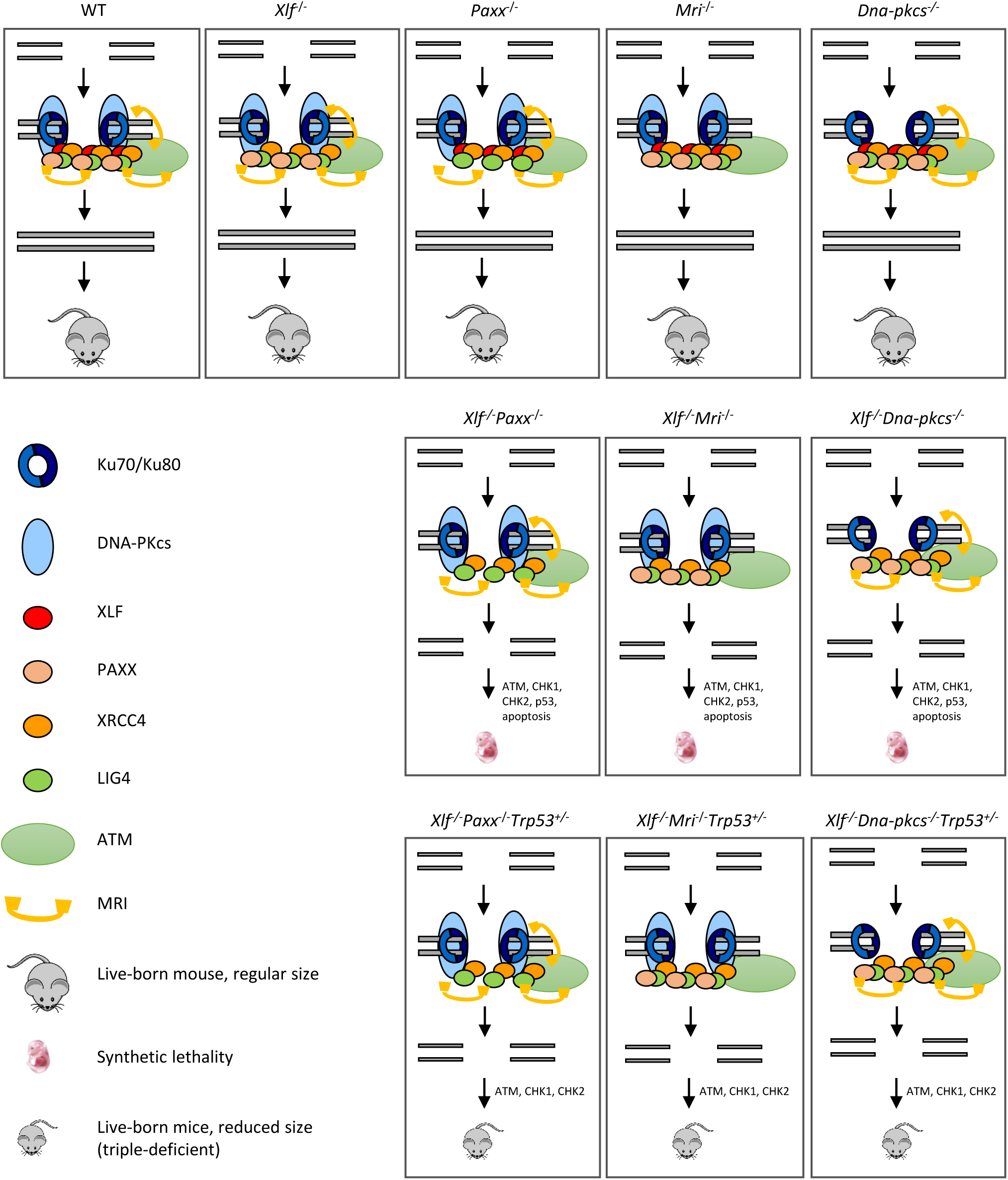
Mutations in NHEJ genes result in distinct phenotypes. Suggested models. Inactivation of *Paxx* or *Mri* results in live-born mice with nearly no DNA repair defects. Inactivation of *Xlf* or *Dna-pkcs* results in live-born mice with increased levels of genomic instability due to reduced NHEJ activity. Combined inactivation of *Xlf/Paxx, Xlf/Mri* or *Xlf/Dna-pkcs* leads to embryonic lethality in mice that correlate with high levels of genomic instability and nearly no NHEJ. Accumulated DSBs activate the ATM-dependent DNA damage response (DDR) pathway; ATM phosphorylates CHK checkpoint proteins that further trigger cell cycle arrest and apoptosis. Alternative end-joining is blocked by presence of Ku70/Ku80. Inactivation of one or two alleles of *Trp53* rescues embryonic lethality of *Xlf/Paxx, Xlf/Mri* and *Xlf/Dna-pkcs* mice. While in these mice the levels of DSBs are increased and ATM-dependent DDR response is activated, lack of p53 prevents massive apoptosis and thus results in alive mice. Sizes of the triple-deficient mice are reduced, as one option, due to DNA damage-dependent cell cycle arrest in multiple cells of the body. The embryonic lethality in mice lacking *Xlf/Paxx* and *Xlf/Mri* is likely to be rescued by inactivation of *Ku70* or *Ku80*.

Our findings have demonstrated that mice lacking XLF, MRI and p53, although live-born, possess a leaky SCID phenotype. *Xlf*^*-/-*^*Mri*^*-/-*^*Trp53*^*+/-*^ mice have a clear fraction of mature B cells in the spleens (CD19+) and bone marrow (B220+CD43-IgM+) (Figures 1 and 2), as well as clear fractions of double- and single-positive T cells in the thymus (CD4+CD8+, CD4+, CD8+) and single-positive T cells in the spleen (CD4+ and CD8+) (Figure 1). However, the cell fractions from these mice are noticeably smaller than those of WT or single-deficient mice. Strikingly, we were able to identify one *Xlf*^*-/-*^*Mri*^*-/-*^ *Trp53*^*+/+*^ mouse at day P30 post-birth. This mouse resembled *Xlf*^*-/-*^*Mri*^*-/-*^*Trp53*^*+/-*^ mice of similar age with respect of B and T cell development, although this mouse was generally sicker than its littermates and had to be euthanized. Similarly, one live-born *Xlf*^*-/-*^*Paxx*^*-/-*^ mouse was reported by *Balmus et al. (2016)* [15], indicating that, exceptionally, embryonic lethality in NHEJ ligation-deficient mice can be overcome, likely due to activity of alternative end-joining. Previously, in 2018, *Hung et al*. [5] reported that combined inactivation of *Xlf* and *Mri* in vAbl pre-B cells results in a severe block in V(D)J recombination and accumulation of unrepaired DSBs *in vitro*, although it was unclear whether this combined inactivation would lead to a deficiency in B lymphocytes when translated to a mouse model [5]. Similarly, double deficient vAbl pre-B cells lacking *Xlf* and *Paxx* are also unable to sustain V(D)J recombination. Importantly, the lack of a progenitor T cell model system left the question of T cell development in *Xlf*^*-/-*^*Mri*^*-/-*^ and *Xlf*^*-/-*^*Paxx*^*-/-*^ mice completely unexplored.

Previously, we showed that mice lacking XLF, PAXX and p53 were live-born and had nearly no B and T cells, reduced size of spleen and hardly detectable thymus [20] (Figure 5). Consistent with this model, a conditional knockout mouse model, which results in double-deficiency of XLF/PAXX in early hematopoietic progenitor cells, was also able to overcome the embryonic lethality of *Xlf*^*-/-*^*Paxx*^*-/-*^ mice [33]. With this model, impairment of V(D)J recombination in *Xlf*^*-/-*^*Paxx*^*-/-*^ cells, as well as the resulting depletion of mature B cells and lack of a visible thymus could also be observed *in vivo* [33]. Our new data provide evidence that *Xlf*^*-/-*^*Paxx*^*-/-*^*Trp53*^*+/-*^ and *Xlf*^*-/-*^*Paxx*^*-/-*^*Trp53*^*-/-*^ mice possess a very small number of mature B cells in the spleen and bone marrow, as well as very minor fractions of single positive T cells in thymus and spleen (Figure 2, 5 and Supplementary Figure 1). Therefore, both mature B and T cells are present in mice lacking XLF/PAXX and XLF/MRI. This can be explained by incomplete blockage in NHEJ and V(D)J recombination, in which the process is dramatically reduced but still possible. We also detected more mature T cells than B cells in these double-deficient mice. Potential explanations include longer lifespan of T cells, which accumulate over time following low efficiency of V(D)J recombination, while B cells are eliminated faster from the pool due to the different physiology [34, 35]. It is also possible that the T cells we detected are a resultant subpopulation that is descendent from the few cells that were able to bypass V(D)J recombination [12]. In this case, the repertoire of T cells based on T cell receptor in mice lacking XLF/PAXX and XLF/MRI would be significantly lower than in control mice, even if normalized to the total cell count. Due to the small presence of mature B and T cells in *Xlf*^*-/-*^*Mri*^*-/-*^*Trp53*^*+/-*^, *Xlf*^*-/-*^*Paxx*^*-/-*^*Trp53*^*+/-*^ and *Xlf*^*-/-*^*Paxx*^*-/-*^*Trp53*^*-/-*^ mice, we categorize the observed immunodeficient phenotypes as “leaky SCID”. Previously, leaky SCID has been described in mice lacking other NHEJ factors, such as *Ku70*^*-/-*^ [6], *Artemis*^*-/-*^ [3], *Lig4*^*-/-*^*Trp53*^*-/-*^ [10, 30], *Xrcc4*^*-/-*^*Trp53*^*-/-*^ [9, 31], *Xlf*^*-/-*^*Atm*^*-/-*^ [19] *and Xlf*^*-/-*^*Rag2*^*c/c*^ [23].

In addition to XLF/MRI and XLF/PAXX deficient mice, inactivation of one or two alleles of *Trp53* also rescues the embryonic lethality of *Xrcc4*^*-/-*^ [9, 31], *Lig4*^*-/-*^ [10, 30] and *Xlf*^*-/-*^*Dna-pkcs*^*-/-*^ [20] mice. We propose a model (Figure 5), when single deficiency for DNA-PKcs, PAXX or MRI results in no or modest phenotypes, and DSBs are efficiently repaired. Combined inactivation of *Xlf/Dna-pkcs, Xlf/Paxx* and *Xlf/Mri* results in inefficient DSB ligation, accumulation of DNA breaks, activation of ATM-dependent DDR, checkpoint protein CHK2, stabilization of p53 and massive apoptosis. This results in embryonic lethality in mice. Furthermore, inactivation of *Trp53* results in *Xlf/Dna-pkcs/Trp53, Xlf/Paxx/Trp53* and *Xlf/Mri/Trp53* triple-deficient mice. While DNA breaks in these mice are not repaired, ATM-dependent DDR response and activation of CHK proteins takes place. However, without p53, apoptosis is not activated, allowing survival of mice (Figure 5). Moreover, we propose that inactivation of *Atm* will also rescue embryonic lethality of *Xlf/Paxx* and *Xlf/Mri* mice due to the mechanisms proposed above. However, inactivation of *Atm* will not rescue embryonic lethality of *Xlf/Dna-pkcs* mice, due to synthetic lethality between *Atm* and *Dna-pkcs*.

It is important to note that altered *Trp53* expression is not always sufficient to rescue embryonic lethality in mice; for example, PLK1-interacting checkpoint helicase (PICH)-deficient mice possess developmental defects in the presence or absence of p53 [36], and ATR mutants (Seckel syndrome) are not completely rescued from embryonic lethality with the inactivation of *Trp53* [37]. Embryonic lethality of XLF/PAXX and XLF/MRI double-deficient mice can be explained by the presence of Ku70/Ku80 heterodimer at the DSBs sites, which blocks DNA repair by alternative end-joining pathway(s), leading to massive apoptosis and cell cycle arrest [38]. Previously, it was shown that embryonic lethality of LIG4-deficient [39] and XLF/DNA-PKcs double-deficient mice [25] could be rescued by inactivating *Ku70* or *Ku80* genes. Similarly, we propose that inactivation of either *Ku70* or *Ku80* gene will rescue the embryonic lethality of XLF/PAXX and XLF/MRI double-deficient mice and will result in mice indistinguishable from Ku70- or Ku80-deficient controls (Figure 5).

Recent studies have shown that *Xlf* genetically interacts with *Rag2* [23] and DDR factors, such as *Atm, 53bp1, H2ax*, and *Mdc1* [17, 19-22, 38]. *Xlf*^*-/-*^*Rag2*^*c/c*^ mice almost completely lack mature B cells and have significantly fewer mature T cells than single deficient controls [23]. *Xlf*^*-/-*^*Atm*^*-/-*^ and *Xlf*^*-/-*^ *53bp1*^*-/-*^ mice are live-born and exhibit reduced body weight, increased genomic instability, and severe lymphocytopenia as a result of V(D)J recombination impairment in developing B and T cells [1, 17, 19, 22]. *Xlf*^*-/-*^*H2ax*^*-/-*^ and *Xlf*^*-/-*^*Mdc1*^*-/-*^, on the other hand, are embryonic lethal [19-21]. There are several possible explanations for the functional redundancy observed between DNA repair genes. For instance, the two factors could have identical (e.g., if both proteins are involved in ligation or DNA end tethering) or complementary (e.g., if one protein stimulates ligation while the other is required for DNA end tethering) functions. To date, XLF has been shown to genetically interact with multiple DNA repair factors [1, 4, 5, 14, 15, 19, 20, 24, 25], and this list is likely to grow [38, 40]. However, no clear genetic interaction has been shown between *Xlf* and *Artemis* or *Xrcc4* in the context of mouse development and V(D)J recombination [24], meaning that it remains difficult to predict genetic interactions without developing and characterizing genetic models.

We found that mice with combined inactivation of *Paxx* and *Mri* (*Paxx*^*-/-*^*Mri*^*-/-*^) are live-born, fertile, and undergo almost normal B and T cell development (Figure 3), where only the number of splenic B cells is affected, giving rise to a modest phenotype. Moreover, inactivation of *Paxx* did not affect the CSR efficiency in *in vitro* stimulated MRI-deficient B cells (Figure 3), thereby confirming our observations *in vitro*. It has been also shown that combined inactivation of *Paxx* and *Mri* genes in vAbl pre-B cells lead to similar V(D)J recombination efficiency to single-deficient *Mri*^*-/-*^, *Paxx*^*-/-*^ and WT controls [5]. Thus, we conclude that there is a genetic interaction between *Paxx* and *Mri*, which results in a modest phenotype.

Lastly, we found that combined inactivation of *Mri* and *Dna-pkcs* (*Mri*^*-/-*^*Dna-pkcs*^*-/-*^) leads to embryonic lethality, and that E14.5 *Mri*^*-/-*^*Dna-pkcs*^*-/-*^ murine embryos were about 40% smaller than single-deficient siblings (Figure 4). DNA-PKcs is associated with the N-terminus of the MRI and Ku heterodimer in the process of recognizing DSBs [5], which may account for genetic interaction between *Mri* and *Dna-pkcs*. Thus, inactivation of *Trp53, Ku70* or *Ku80* may be a viable method to rescue synthetic lethality from *Mri*^*-/-*^*Dna-pkcs*^*-/-*^ mice.

In conclusion, we have developed and described several complex genetic mouse models (Figure 5). *Xlf*^*-/-*^*Mri*^*-/-*^*Trp53*^*+/-*^ and *Xlf*^*-/-*^*Paxx*^*-/-*^*Trp53*^*+(-)/-*^ mice possessed severely impaired B and T lymphocyte development, leaky SCID; *Paxx*^*-/-*^*Mri*^*-/-*^ mice develop a modest B cell phenotype; and *Mri*^*-/-*^ *Dna-pkcs*^*-/-*^ mice are embryonic lethal.

## Author contributions

VO, SCZ, QZ, AL and MFB designed the study, analyzed and interpreted the results. SCZ, QZ, AL and MFB performed most of the experiments. VO wrote the paper with the help of SCZ and RY. All the authors contributed to writing of the final manuscript.

## Acknowledgments

This work was supported by the Research Council of Norway Young Talent Investigator grant (#249774) to V.O. In addition, VO group was supported by the Liaison Committee for Education, Research, and Innovation in Central Norway (#13477; #38811); the Norwegian Cancer Society (#182355); the Research Council of Norway FRIMEDBIO grants (#270491 and #291217), and The Outstanding Academic Fellow Program at NTNU (2017–2021).

---

**Supplementary Figure 1.**
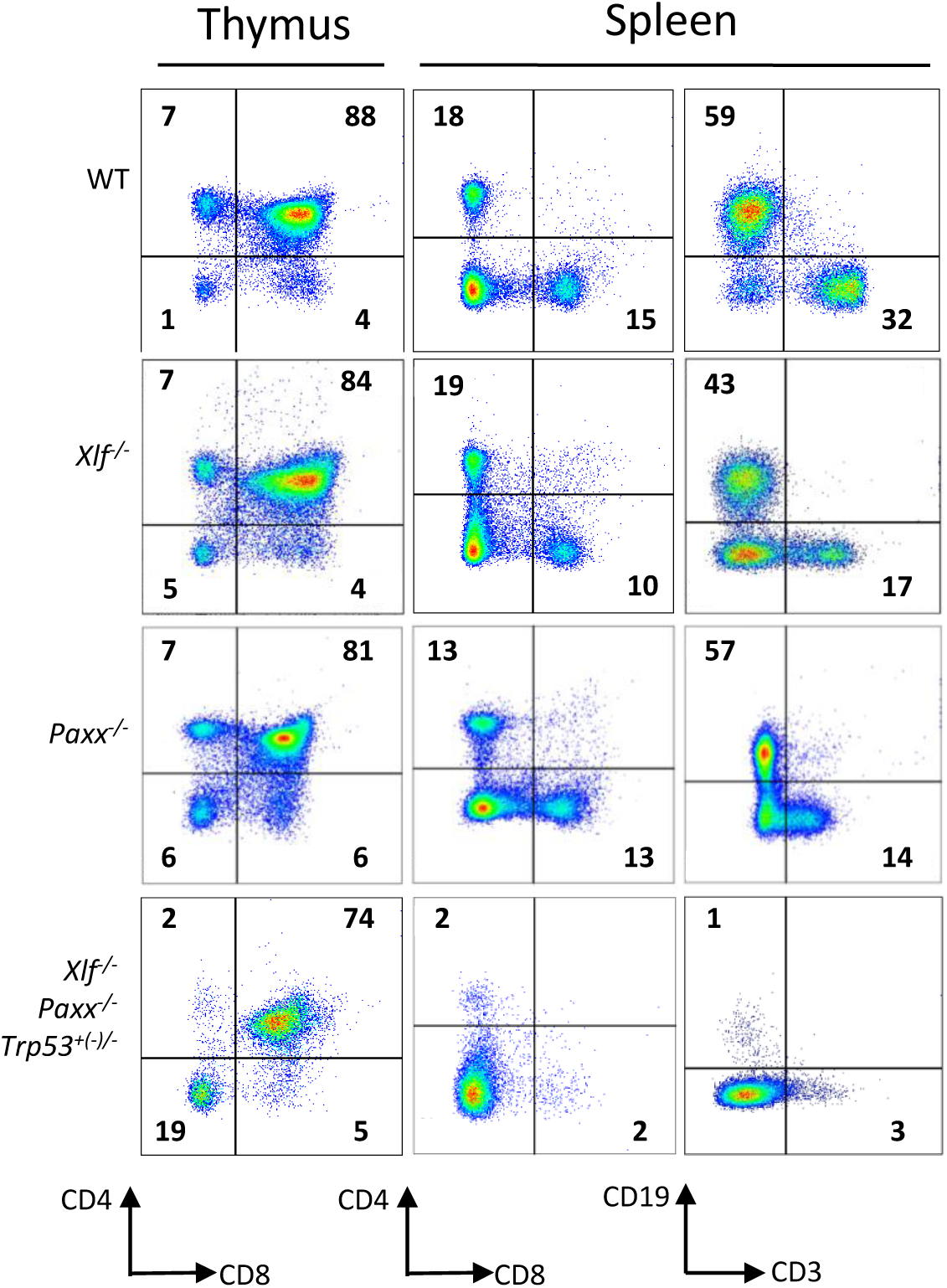
Development of B and T cells in *Xlf*^*-/-*^*Paxx*^*-/-*^*Trp53*^*+(-)/-*^ mice. Examples of flow cytometric analysis of thymic and splenic T cell subsets and splenic CD19+ B cells. *Xlf*^*-/-*^*Paxx*^*-/-*^*Trp53*^*+(-)/-*^ is a combination of *Xlf*^*-/-*^*Paxx*^*-/-*^*Trp53*^*+/-*^ and *Xlf*^*-/-*^*Paxx*^*-/-*^*Trp53*^*-/-*^.

## References

1. Kumar V, Alt FW, Oksenych V. Functional overlaps between XLF and the ATM-dependent DNA double strand break response. DNA Repair (Amst). 2014; 16: 11–22. https://doi.org:10.1016/j.dnarep.2014.01.010

2. Gao Y, Chaudhuri J, Zhu C, Davidson L, Weaver DT, Alt FW. A targeted DNA-PKcs-null mutation reveals DNA-PK-independent functions for KU in V(D)J recombination. Immunity. 1998; 9: 367–76. https://doi.org:10.1016/s1074-7613(00)80619-6

3. Rooney S, Sekiguchi J, Zhu C, Cheng HL, Manis J, Whitlow S, DeVido J, Foy D, Chaudhuri J, Lombard D, Alt FW. Leaky Scid phenotype associated with defective V(D)J coding end processing in Artemis-deficient mice. Mol Cell. 2002; 10: 1379–90. https://doi.org:10.1016/s1097-2765(02)00755-4

4. Liu X, Shao Z, Jiang W, Lee BJ, Zha S. PAXX promotes KU accumulation at DNA breaks and is essential for end-joining in XLF-deficient mice. Nat Commun. 2017; 8: 13816. https://doi.org:10.1038/ncomms13816

5. Hung PJ, Johnson B, Chen BR, Byrum AK, Bredemeyer AL, Yewdell WT, Johnson TE, Lee BJ, Deivasigamani S, Hindi I, Amatya P, Gross ML, Paull TT, et al. MRI Is a DNA Damage Response Adaptor during Classical Non-homologous End Joining. Mol Cell. 2018; 71: 332–42 e8. https://doi.org:10.1016/j.molcel.2018.06.018

6. Gu Y, Seidl KJ, Rathbun GA, Zhu C, Manis JP, van der Stoep N, Davidson L, Cheng HL, Sekiguchi JM, Frank K, Stanhope-Baker P, Schlissel MS, Roth DB, et al. Growth retardation and leaky SCID phenotype of Ku70-deficient mice. Immunity. 1997; 7: 653–65. https://doi.org:10.1016/S1074-7613(00)80386-6

7. Nussenzweig A, Chen C, da Costa Soares V, Sanchez M, Sokol K, Nussenzweig MC, Li GC. Requirement for Ku80 in growth and immunoglobulin V(D)J recombination. Nature. 1996; 382: 551–5. https://doi.org:10.1038/382551a0

8. Ma Y, Pannicke U, Schwarz K, Lieber MR. Hairpin opening and overhang processing by an Artemis/DNA-dependent protein kinase complex in nonhomologous end joining and V(D)J recombination. Cell. 2002; 108: 781–94. https://doi.org:10.1016/s0092-8674(02)00671-2

9. Gao Y, Sun Y, Frank KM, Dikkes P, Fujiwara Y, Seidl KJ, Sekiguchi JM, Rathbun GA, Swat W, Wang J, Bronson RT, Malynn BA, Bryans M, et al. A critical role for DNA end-joining proteins in both lymphogenesis and neurogenesis. Cell. 1998; 95: 891–902.

10. Frank KM, Sekiguchi JM, Seidl KJ, Swat W, Rathbun GA, Cheng HL, Davidson L, Kangaloo L, Alt FW. Late embryonic lethality and impaired V(D)J recombination in mice lacking DNA ligase IV. Nature. 1998; 396: 173–7. https://doi.org:10.1038/24172

11. Li G, Alt FW, Cheng HL, Brush JW, Goff PH, Murphy MM, Franco S, Zhang Y, Zha S. Lymphocyte-specific compensation for XLF/cernunnos end-joining functions in V(D)J recombination. Mol Cell. 2008; 31: 631–40. https://doi.org:10.1016/j.molcel.2008.07.017

12. Vera G, Rivera-Munoz P, Abramowski V, Malivert L, Lim A, Bole-Feysot C, Martin C, Florkin B, Latour S, Revy P, de Villartay JP. Cernunnos deficiency reduces thymocyte life span and alters the T cell repertoire in mice and humans. Mol Cell Biol. 2013; 33: 701–11. https://doi.org:10.1128/MCB.01057-12

13. Roch B, Abramowski V, Chaumeil J, de Villartay JP. Cernunnos/Xlf Deficiency Results in Suboptimal V(D)J Recombination and Impaired Lymphoid Development in Mice. Front Immunol. 2019; 10: 443. https://doi.org:10.3389/fimmu.2019.00443

14. Abramowski V, Etienne O, Elsaid R, Yang J, Berland A, Kermasson L, Roch B, Musilli S, Moussu JP, Lipson-Ruffert K, Revy P, Cumano A, Boussin FD, et al. PAXX and Xlf interplay revealed by impaired CNS development and immunodeficiency of double KO mice. Cell Death Differ. 2018; 25: 444–52. https://doi.org:10.1038/cdd.2017.184

15. Balmus G, Barros AC, Wijnhoven PW, Lescale C, Hasse HL, Boroviak K, le Sage C, Doe B, Speak AO, Galli A, Jacobsen M, Deriano L, Adams DJ, et al. Synthetic lethality between PAXX and XLF in mammalian development. Genes Dev. 2016; 30: 2152–7. https://doi.org:10.1101/gad.290510.116

16. Gago-Fuentes R, Xing M, Saeterstad S, Sarno A, Dewan A, Beck C, Bradamante S, Bjoras M, Oksenych V. Normal development of mice lacking PAXX, the paralogue of XRCC4 and XLF. FEBS Open Bio. 2018; 8: 426–34. https://doi.org:10.1002/2211-5463.12381

17. Liu X, Jiang W, Dubois RL, Yamamoto K, Wolner Z, Zha S. Overlapping functions between XLF repair protein and 53BP1 DNA damage response factor in end joining and lymphocyte development. Proc Natl Acad Sci U S A. 2012; 109: 3903–8. https://doi.org:10.1073/pnas.1120160109

18. Castaneda-Zegarra S, Huse C, Røsand Ø, Sarno A, Xing M, Gago-Fuentes R, Zhang Q, Alirezaylavasani A, Werner J, Ji P, Liabakk N, Wang W, Bjørås M, et al. Generation of a Mouse Model Lacking the Non-Homologous End-Joining Factor Mri/Cyren. Biomolecules. 2019; 9. https://doi.org:10.3390/biom9120798

19. Zha S, Guo C, Boboila C, Oksenych V, Cheng HL, Zhang Y, Wesemann DR, Yuen G, Patel H, Goff PH, Dubois RL, Alt FW. ATM damage response and XLF repair factor are functionally redundant in joining DNA breaks. Nature. 2011; 469: 250–4. https://doi.org:10.1038/nature09604

20. Castaneda-Zegarra S, Xing M, Gago-Fuentes R, Saeterstad S, Oksenych V. Synthetic lethality between DNA repair factors Xlf and Paxx is rescued by inactivation of Trp53. DNA Repair (Amst). 2019; 73: 164–9. https://doi.org:10.1016/j.dnarep.2018.12.002

21. Beck C, Castaneda-Zegarra S, Huse C, Xing M, Oksenych V. Mediator of DNA Damage Checkpoint Protein 1 Facilitates V(D)J Recombination in Cells Lacking DNA Repair Factor XLF. Biomolecules. 2019; 10. https://doi.org:10.3390/biom10010060

22. Oksenych V, Alt FW, Kumar V, Schwer B, Wesemann DR, Hansen E, Patel H, Su A, Guo C. Functional redundancy between repair factor XLF and damage response mediator 53BP1 in V(D)J recombination and DNA repair. Proc Natl Acad Sci U S A. 2012; 109: 2455–60. https://doi.org:10.1073/pnas.1121458109

23. Lescale C, Abramowski V, Bedora-Faure M, Murigneux V, Vera G, Roth DB, Revy P, de Villartay JP, Deriano L. RAG2 and XLF/Cernunnos interplay reveals a novel role for the RAG complex in DNA repair. Nat Commun. 2016; 7: 10529. https://doi.org:10.1038/ncomms10529

24. Oksenych V, Kumar V, Liu X, Guo C, Schwer B, Zha S, Alt FW. Functional redundancy between the XLF and DNA-PKcs DNA repair factors in V(D)J recombination and nonhomologous DNA end joining. Proc Natl Acad Sci U S A. 2013; 110: 2234–9. https://doi.org:10.1073/pnas.1222573110

25. Xing M, Bjoras M, Daniel JA, Alt FW, Oksenych V. Synthetic lethality between murine DNA repair factors XLF and DNA-PKcs is rescued by inactivation of Ku70. DNA Repair (Amst). 2017; 57: 133–8. https://doi.org:10.1016/j.dnarep.2017.07.008

26. Lescale C, Lenden Hasse H, Blackford AN, Balmus G, Bianchi JJ, Yu W, Bacoccina L, Jarade A, Clouin C, Sivapalan R, Reina-San-Martin B, Jackson SP, Deriano L. Specific Roles of XRCC4 Paralogs PAXX and XLF during V(D)J Recombination. Cell Rep. 2016; 16: 2967–79. https://doi.org:10.1016/j.celrep.2016.08.069

27. Kumar V, Alt FW, Frock RL. PAXX and XLF DNA repair factors are functionally redundant in joining DNA breaks in a G1-arrested progenitor B-cell line. Proc Natl Acad Sci U S A. 2016; 113: 10619–24. https://doi.org:10.1073/pnas.1611882113

28. Hung PJ, Chen BR, George R, Liberman C, Morales AJ, Colon-Ortiz P, Tyler JK, Sleckman BP, Bredemeyer AL. Deficiency of XLF and PAXX prevents DNA double-strand break repair by non-homologous end joining in lymphocytes. Cell Cycle. 2017; 16: 286–95. https://doi.org:10.1080/15384101.2016.1253640

29. Barnes DE, Stamp G, Rosewell I, Denzel A, Lindahl T. Targeted disruption of the gene encoding DNA ligase IV leads to lethality in embryonic mice. Curr Biol. 1998; 8: 1395–8. https://doi.org:10.1016/s0960-9822(98)00021-9

30. Frank KM, Sharpless NE, Gao Y, Sekiguchi JM, Ferguson DO, Zhu C, Manis JP, Horner J, DePinho RA, Alt FW. DNA ligase IV deficiency in mice leads to defective neurogenesis and embryonic lethality via the p53 pathway. Mol Cell. 2000; 5: 993–1002.

31. Gao Y, Ferguson DO, Xie W, Manis JP, Sekiguchi J, Frank KM, Chaudhuri J, Horner J, DePinho RA, Alt FW. Interplay of p53 and DNA-repair protein XRCC4 in tumorigenesis, genomic stability and development. Nature. 2000; 404: 897–900. https://doi.org:10.1038/35009138

32. Jacks T, Remington L, Williams BO, Schmitt EM, Halachmi S, Bronson RT, Weinberg RA. Tumor spectrum analysis in p53-mutant mice. Curr Biol. 1994; 4: 1–7. https://doi.org:10.1016/s0960-9822(00)00002-6

33. Musilli S, Abramowski V, Roch B, de Villartay JP. An in vivo study of the impact of deficiency in the DNA repair proteins PAXX and XLF on development and maturation of the hemolymphoid system. J Biol Chem. 2020; 295: 2398–406. https://doi.org:10.1074/jbc.AC119.010924

34. Di Rosa F, Ramaswamy S, Ridge JP, Matzinger P. On the lifespan of virgin T lymphocytes. J Immunol. 1999; 163: 1253–7.

35. Fulcher DA, Basten A. B cell life span: a review. Immunol Cell Biol. 1997; 75: 446–55. https://doi.org:10.1038/icb.1997.69

36. Albers E, Sbroggio M, Pladevall-Morera D, Bizard AH, Avram A, Gonzalez P, Martin-Gonzalez J, Hickson ID, Lopez-Contreras AJ. Loss of PICH Results in Chromosomal Instability, p53 Activation, and Embryonic Lethality. Cell Rep. 2018; 24: 3274–84. https://doi.org:10.1016/j.celrep.2018.08.071

37. Murga M, Bunting S, Montana MF, Soria R, Mulero F, Canamero M, Lee Y, McKinnon PJ, Nussenzweig A, Fernandez-Capetillo O. A mouse model of ATR-Seckel shows embryonic replicative stress and accelerated aging. Nat Genet. 2009; 41: 891–8. https://doi.org:10.1038/ng.420

38. Castaneda-Zegarra S, Fernandez-Berrocal M, Tkachev M, Yao R, Upfold NLE, Oksenych V. Genetic interaction between the non-homologous end joining factors during B and T lymphocyte development: in vivo mouse models. Scand J Immunol. 2020: e12936. https://doi.org:10.1111/sji.12936

39. Karanjawala ZE, Adachi N, Irvine RA, Oh EK, Shibata D, Schwarz K, Hsieh CL, Lieber MR. The embryonic lethality in DNA ligase IV-deficient mice is rescued by deletion of Ku: implications for unifying the heterogeneous phenotypes of NHEJ mutants. DNA Repair (Amst). 2002; 1: 1017–26. https://doi.org:10.1016/s1568-7864(02)00151-9

40. Wang X, Lee B, Zha S. The recent advances in non-homologous end-joining through the lens of lymphocyte development. DNA Repair (Amst). 2020. https://doi.org:10.1016/j.dnarep.2020.102874

